# Evolutionary Adaptations of IRG1 Refines Itaconate Synthesis and Mitigates Innate Immunometabolism Trade-offs

**DOI:** 10.1101/2022.06.17.496652

**Authors:** Richard V. Szeligowski, Francois Miros, Andres Saez, Marisa DeCiucis, Gunter P. Wagner, Hongying Shen

## Abstract

Itaconate is an innate immune metabolite specifically produced in activated immune cells via the decarboxylation of *cis*-aconitate, an intermediate of the TCA cycle. By inhibiting succinate-related metabolism, itaconate exerts antimicrobial properties at the expense of potentially disrupting the hosts’ own central energy metabolism, a double-edged dilemma of immunometabolism. To explore the evolutionary logic of itaconate biosynthesis, we investigated the evolutionary trajectory of IRG1, which encodes for *cis*-aconitate decarboxylase (CAD), the enzyme responsible for itaconate production. Phylogenetic analysis reveals a putative independent acquisition of metazoan and fungal IRG1 from prokaryotic sources. In metazoans, IRG1 underwent gene duplication and subsequently lost the mitochondrial targeting sequence (MTS), relocating CAD outside the mitochondrial matrix and therefore preventing direct inhibition of energy metabolism. In basal metazoans that contain IRG1, oysters and amphioxus, primitive IRG1 expression is also induced by innate immune stimuli, suggesting an already specialized role of itaconate for innate immune defense in early bilaterians. Our integrated *in silico* and experimental analysis highlight the molecular adaptations in IRG1, including subcellular relocation, that optimize itaconate production for innate immunity in resolving a fundamental trade-off in immunometabolism.

## Introduction

Recent multispecies analysis reveals that enzyme biochemistry dedicated to modern eukaryotic innate immunity has deep evolutionary roots, with a specialized role in the innate immunity of ancient life. Examples include the cGAS-STING pathway in animals that shares ancestry with the prokaryotic cyclic-oligonucleotide in anti-phage defense^1–4^, and the eukaryotic peptidoglycan amidase that digests the bacterial cell wall and was horizontally acquired from prokaryotic antibacterial toxins^5^. We rationalize that tracing the evolutionary trajectory of the specialized biochemistry in innate immunity might provide insights into the regulatory mechanisms that shape and optimize the modern innate immune response compatible with the organismal fitness.

Itaconate, an innate immunometabolite with both antimicrobial and anti-inflammatory roles^6–8^ is an exemplar of specialized biochemistry in mammalian innate immunity. Itaconate is produced via a single decarboxylation step by the enzyme *cis*-aconitate decarboxylase, the gene product of the immune response gene (IRG1, also named ACOD1)^9^, shunting away the central energy metabolic pathway TCA cycle intermediate *cis*-aconitate at a high flux rate (Fig. 1A). Itaconate is highly reactive due to the terminal double bond, the alkene group on the C2 position (Fig. 1A). This reactivity enables antimicrobial activity via inhibiting isocitrate lyase in the glyoxylate shunt, a key energy metabolic pathway exclusively found in prokaryotes^9–11^, and vitamin B12-dependent methylmalonyl-CoA mutase-mediated odd chain carbon utilization^12,13^. However, in the host, itaconate also inhibits key mitochondrial matrix-localized energy metabolism including succinyl-CoA ligase^14^, succinate dehydrogenase^15,16^ and B12-dependent methylmalonyl-CoA mutase producing succinate^12^, in addition to functioning as a Michael acceptor reacting to nucleophilic sulfur group of the cysteine residue of KEAP1 activating NRF2^17^ and that of the antioxidant glutathione^18^. Itaconate can be completely degraded by a citramalyl-CoA lyase activity (Fig. 1A) that was discovered for human CLYBL gene protecting the vitamin B12 cofactor^12^. The same activity of bacterial citE provided fitness (virulence) to the bacterial pathogen by degrading itaconate to counter the host defense^19^, which illustrates an evolutionary arms race.

**Figure 1.**
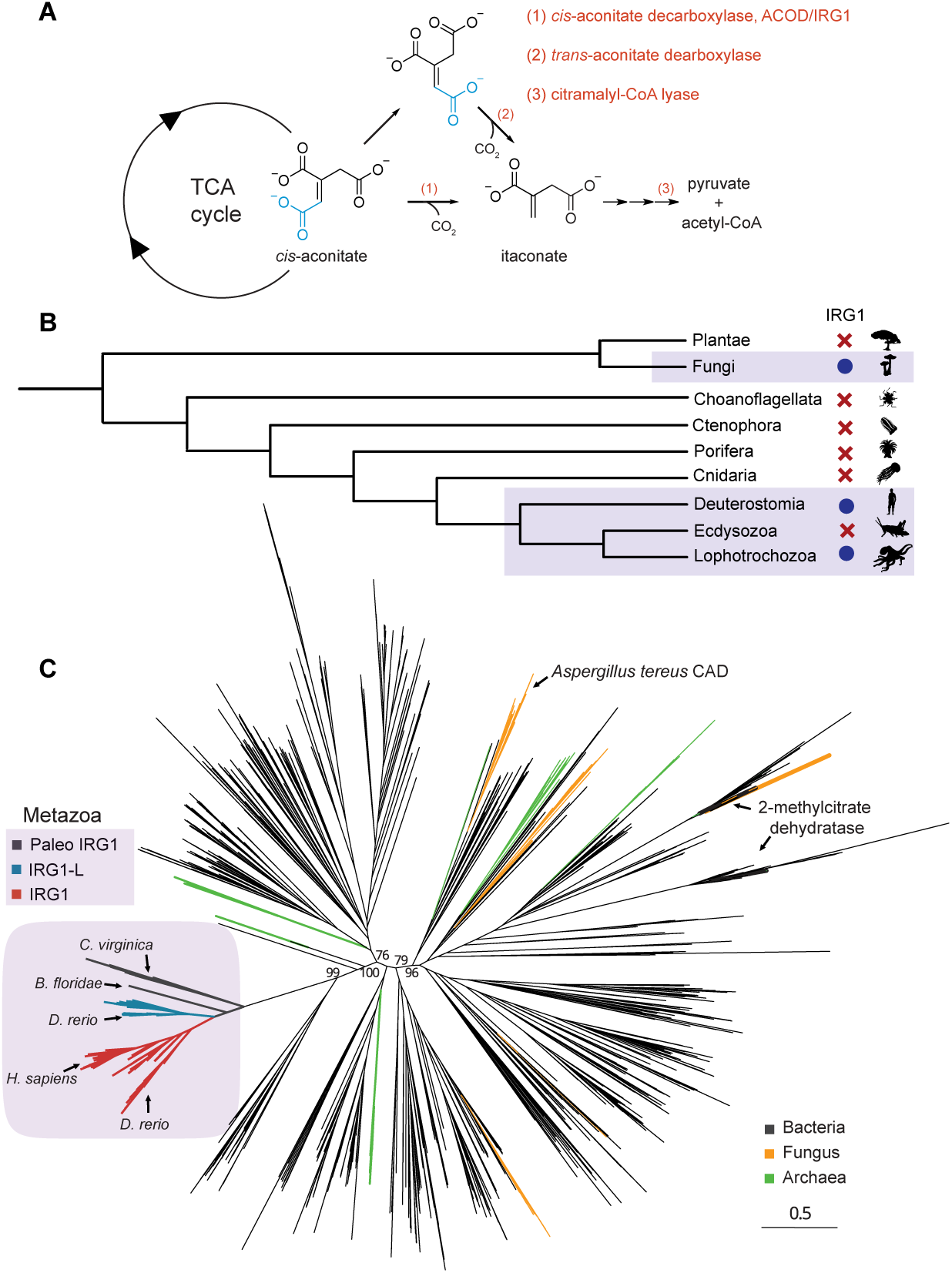
Taxonomic distribution and molecular phylogeny of IRG1, the enzyme producing the innate immune metabolite itaconate. (A) Itaconate biosynthesis and degradation pathways in nature. Two itaconate biosynthesis pathways produce itaconate from the TCA cycle intermediate *cis*-aconitate, through either the *cis*-aconitate decarboxylase (CAD) enzyme IRG1/ACOD1 or the trans-aconitate decarboxylase (TAD) enzyme. Itaconate degradation involves a dedicated citramalyl-CoA lyase CLYBL to generate pyruvate and acetyl-CoA. (B) Taxonomic distribution of IRG1 in eukaryotes. IRG1 is present in Fungi, Lophotrochozoans, and Deuterostomes but is absent in other metazoans. (C) A phylogenetic tree of IRG1 homologs in nature. Bacterial sequences are colored in black, archaeal in green and fungal in orange. High confidence bootstrapping values are shown for the main branches separating the metazoan and *A.tereus* CAD branches. Branches containing experimentally validated 2-methylcitrate dehydratases are labeled. In the Metazoan branch, sequences were colored based on three lineages, IRG1/ACOD1 (red), IRG1L (IRG1-like, cyan) and Paleo-IRG1 (pIRG1, black). Metazoan proteins that are subsequently studied here are labeled, *C. virginica* (eastern oyster) pIRG1, *B. floridae* (lancelet) pIRG1 and *D.rerio* (zebrafish) IRG1L, in comparison to *D.rerio* (zebrafish) IRG1/ACOD1 and *H.sapiens* (human) IRG1/ACOD1.

Taken together, itaconate’s dual inhibition of the host and pathogen’s central energy metabolic pathway presents a “double-edged” dilemma in immunometabolism: the host benefits from the elimination of pathogens, but also suffers an apparent detrimental effect via the disruption of the organism’s own energy metabolism.

One mechanism to alleviate this dilemma involves a spatiotemporal restricted transcriptional regulation of IRG1, where its expression is exclusively restricted to macrophages in humans, mice, and zebrafish^20^. Within macrophages, IRG1 transcription is strictly kept silent and only upregulated upon innate immune stimulation by bacterial pathogens or LPS. Only a few studies reported the induction of IRG1 in other cell types by other stimuli, including IRG1 paralog in zebrafish that is induced in terminally differentiated keratinocytes upon bacterial infection^21^ and the induction of human IRG1 in terminally differentiated neurons upon zika virus infection^22^.

This highly selective transcriptional hierarchy for IRG1 is evidently successful in resolving the “double-edged” dilemma in modern vertebrates. However, other mechanisms to resolve this dilemma may have arisen in distant species where transcriptional regulation is less sophisticated.

To explore the evolutionary logic of itaconate production, we investigated the taxonomic distribution and the evolutionary trajectory of IRG1 *in silico*. We discovered that fungal IRG1 and metazoan IRG1 were likely independently acquired from prokaryotes. Whole genome duplication early in vertebrate history, specifically in the gnathstome stem lineage, produced two paralogous IRG1 genes and subsequent loss events in therian mammal lineages (marsupials and placental mammals) enabled a precise regulation of itaconate biosynthesis by removing the mitochondrial targeting sequence (MTS) in IRG1 that prevents a direct inhibition of mitochondrial matrix bioenergetics. Our cell- and tissue-based studies in the most distantly diverged metazoan species that contain IRG1 (paleo-IRG1), mollusks (eastern oyster, *Crassostrea virginica*) and basal chordates (amphioxus, *Branchiostoma floridae*), identified a transcriptional induction of IRG1 by innate immune inducers, suggesting an ancestrally dedicated innate immune role for IRG1. Together, our molecular phylogeny and non-model organism studies reveal a multilayered evolutionary strategy that tightens the regulations of itaconate production. We speculate that similar evolutionary regulatory mechanisms may be broadly utilized to shape toxic innate immune biochemistry and to resolve the “double-edged” dilemma in immune defense.

## Results

To understand the occurrence of itaconate production in nature, our initial literature survey identified two enzymes that synthesize itaconate: the cis-aconitate decarboxylase (CAD) enzyme in humans, mice^9^, zebrafish^20^, and the fungus *Aspergillus terreus*^23,24^, and *trans*-aconitate decarboxylase (TAD) in *Ustilago maydis*, a plant pathogenic fungus^25^ (Fig. 1A). Primary sequence analysis and domain searches suggest that all CAD sequences are comprised of the MmgE/PrpD domain fold^26^, a protein domain family that also contains the prokaryotic methylcitrate dehydratase in the methylcitrate cycle^27^. The TAD enzymes are comprised of a completely different protein fold that is shared by the adenylosuccinate lyase enzyme ADSL in mammalian purine nucleotide biosynthesis. However, TAD enzymes appear to be exclusively restricted to the fungi kingdom. For CAD, many fungal studies focus on the bioengineering role of itaconate production for polymer biosynthesis^28^, and the physiological role of *A. terreus* CAD and *U. maydis* TAD is poorly understood — a potential function in immune defense may hint at convergent evolution to produce itaconate starting from different protein folds^29^.

Because CAD enzymes exhibit a broad taxonomic distribution spanning fungi and mammals, we then focused on the evolutionary trajectory of CAD across the tree of life. Combined BLAST searches (BLASTP and TBLASTN searches using human IRG1 protein sequence as a query) and a MmgE/PrpD Pfam domain search returned protein sequences spanning all three domains of life in bacteria, archaea and eukaryotes. A closer inspection of the eukaryotic domain suggests a sporadic presence of IRG1 homologs in fungi, deuterostomes, and lophotrochozoans, but CAD is notably absent in their sister clades including ecdysozoans (arthropods, nematodes and their relatives) and plants (Fig. 1B). Six putative Drosophila sequences were determined to be misannotated and thus excluded for further analysis (see methods). The MmgE/PrpD domain-containing proteins are present in Basidiomycota and Ascomycota, the fungi taxa including *A. terreus*, but are absent in other basal fungi taxa including Chytridiomycota and Zygomycota. The patchiness of IRG1 occurrence in nature suggests that itaconate biosynthesis is not critical, and thus dispensable for eukaryotic central energy metabolism and organismal fitness.

Molecular phylogenetic analysis of all prokaryotic and eukaryotic IRG1 homologs (Supplementary File 1) highlighted a clear separation of the fungi and animal IRG1 sequences, suggesting two independent acquisition events. We first confirmed that all the IRG1 sequences contain the MmgE/PrpD domain and then manually added eight experimentally validated 2-methylcitrate dehydratases including bacterial sequences from *E. coli*^30^, *S. typhimurium*^31^, *C. necator*^32^*, C. glutamicum*^33^*, B. subtilis*^34^, and two different homologs in *M. tuberculosis*^35–37^ as well as a fungal sequence from *Y. lipolytica*^38^. The most striking observation from the phylogenetic analysis is that, despite similar enzymatic activities, the validated fungi *A. terreus* CAD and the metazoan IRG1 sequences reside on two distant branches intercalated by prokaryotic sequences with high bootstrapping values (>75) (Fig. 1C), strongly suggesting that metazoan CAD/ IRG1 and the fungi CAD are independently acquired.

The absence of the CAD enzyme sequences in many fungal taxa and the close relationship of *A. terreus* CAD sequence with the prokaryotic sequences in adjacent branches may indicate a putative direct horizontal gene transfer (HGT) into the *Aspergillus* ancestor. The eight experimentally validated 2-methylcitrate dehydratases are clustered on a separate branch from both *A. terreus* CAD and metazoan IRG1, suggesting they are indeed different enzymes that can be distinguished by sequence homology. Notably, the nearest clade to metazoan IRG1 orthologs contain alphaproteobacterial sequences including one from the Rickettsiales genera, the bacterial taxa considered to be the closest extant organism to eukaryotic mitochondria^39^. However, the origin of prokaryotic clades where metazoans acquired IRG1 remains unknown.

The evolutionary trajectory of metazoan IRG1 is consistent with canonical metazoan phylogeny and is composed of three distinct subbranches: a homolog in mollusks and basal chordates we called paleo-IRG1 (pIRG1, Fig. 2, in gray) and two paralogous vertebrate groups that we designated IRG1L (IRG1-like, Fig. 2, in blue) and IRG1 (Fig. 2, in red). The pIRG1 sequences consist of invertebrate sequences exclusively in mollusks, the basal chordate amphioxus (*Branchiostoma* sp.), and agnathans (lamprey, *Petromyzon marinus*) (Fig. 1C). A clear divergence of the IRG1L and IRG1 branches in several gnathostome classes (in Actinopterygii, Sarcopterygii, Amphibia, Aves, and Reptilia) and the conspicuous absence of IRG1L in therian mammals prompted us to further inspect the taxonomic distribution and phylogeny of IRG1s in metazoans.

**Figure 2.**
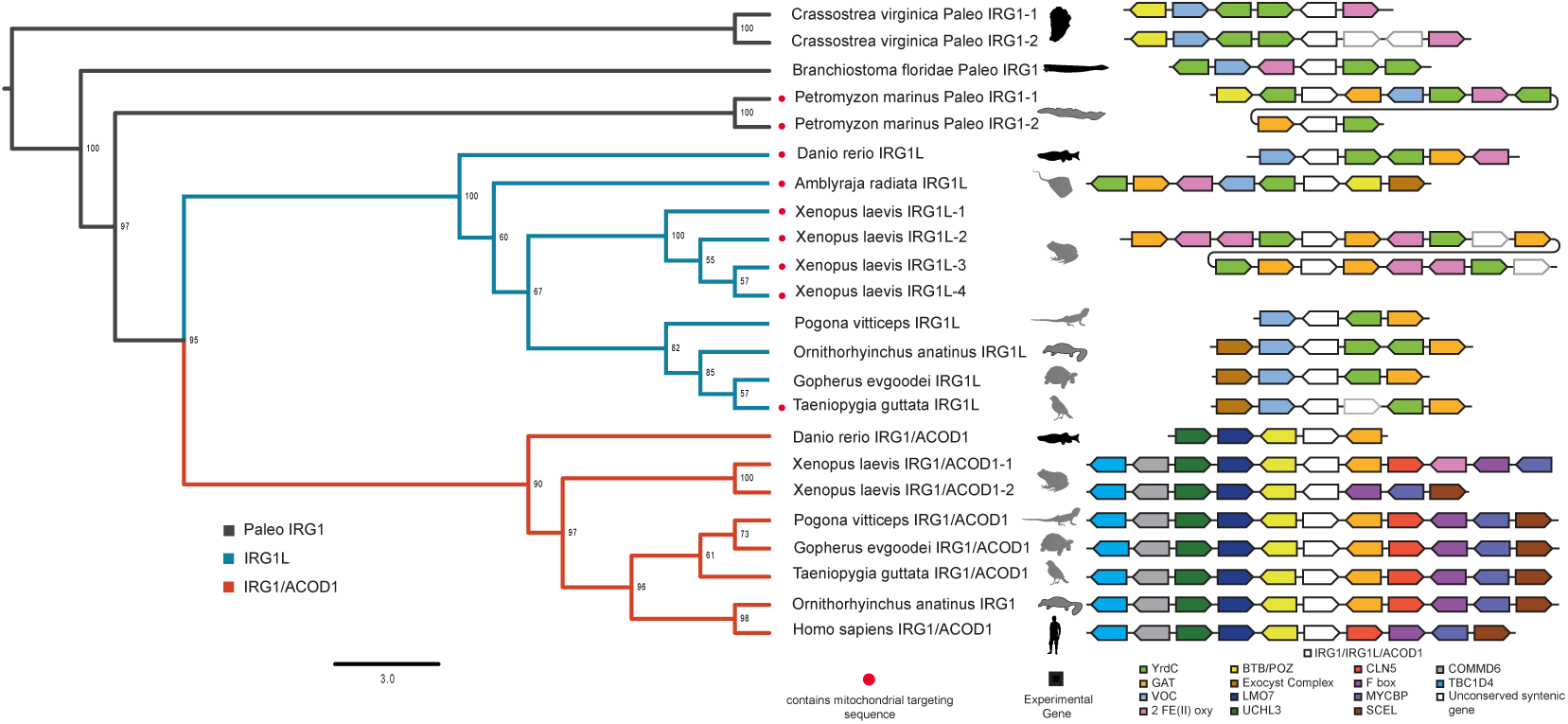
Phylogeny and synteny for representative metazoan IRG1 sequences. The tree branches are color-coded for the three lineages including paleo-IRG1 (p IRG1, black), IRG1-like (IRG1L, cyan) and IRG1 (red). The presence of the predicted Mitochondrial Targeting Sequence (MTS) is highlighted by the red dots. Homologous genes in the genomic vicinity spanning the IRG1, IRG1L and IRG1 are color-coded, for which the homologs present in human were annotated by human gene names, or otherwise by Pfam protein domain name. The two copies of pIRG1 in *C. virginica* are separated by approximately 200 kb, indicated by a dashed line in the syntenic block. Images for experimentally examined species in this report are shown in black, including easter oyster (*C. virginica*), amphioxus (*B. floridae*), and zebrafish (*D. rerio*) IRG1L.

We then focused on the metazoan tree and color-coded the sequences in the three subbranches corresponding to pIRG1, IRG1 and IRG1L, by taxonomy (Fig. S1A). After analyzing the taxonomic distribution of the IRG1 homologs, we confirmed that all IRG1 and IRG1L sequences belong to jawed vertebrates and that each paralogous branch follows a canonical vertebrate phylogeny. The presence of both IRG1 and IRG1L in the majority of jawed vertebrate taxa, including bony fish, amphibians, reptiles, and birds and the exclusive presence of pIRG1 in mollusks and agnathans strongly support that IRG1 and IRG1L are derived from the pIRG1 through the putative genome duplication event in the gnathostome stem-lineage^40^.

Several taxon-specific local gene duplication events were observed, including two syntenic pIRG1s in oysters (> 99.48% nucleotide sequence identity), two syntenic pIRG1s in lamprey (> 98.72% nucleotide sequence identity), four syntenic IRG1Ls in *Xenopus laevis* (Fig. 2 and Fig. S1), and two IRG1Ls in salmonid fish likely due to known genome duplication in these taxa^41^.

Cartilaginous fish genomes were found to only contain IRG1L, but lack IRG1 (Fig. S1A), a result we further confirmed by a manual BLAST search in species including the skate (*Amblyraja radiata*) and sharks (*Chilscyllium plagiosum, Rhincodon typus, Chiloscyllium punctatum, Carcharodon carcharias, Scyliorhinus torazame, Callorhinchus milii and Scyliorhinus canicula*).

The most striking observation in the taxa-annotated metazoan tree is a complete loss of IRG1L in the metatherian and eutherian mammals that otherwise exhibit an expansion of IRG1 lineage (Fig. S1A, pink), which contrasts with the presence of both IRG1 and IRG1L in the prototherian platypus (*O. anatinus*; Figure S1A, arrows). To further confirm the difference, we manually searched for IRG1 and IRG1L in major mammalian orders^42^ by either BlastP or TblastN search using human IRG1 and zebrafish IRG1L sequences (Fig. S1B). The absence of both IRG1 and IRG1L in Hyracoidea and Xenarthra (Fig. S1B) is consistent with the dispensability of IRG1 for organismal energy metabolism. Thus, all metatherians and eutherians contain only IRG1. Within the mammalian clade, IRG1L is exclusive to prototheria, a clade of egg-laying mammals, and is completely lost in metatheria and eutheria, the viviparous mammals (Fig. S1B), inviting further analysis in the difference between IRG1L and IRG1.

Primary sequence inspection unexpectedly revealed a N-terminally extended amino acid sequence that is frequently present in both IRG1L and pIRG1 sequences, but much less frequent in the IRG1 sequences. Because the CAD enzyme directly acts on intermediates of the mitochondrial matrix-localized TCA cycle, we then explored if these N-terminal extensions might encode the canonical mitochondrial targeting sequence (MTS)^43^, the peptide motif directing proteins into the mitochondrial matrix. Indeed, investigating all metazoan sequences using MTS prediction software TargetP revealed high-confidence MTS present in 34.56% pIRG1 and IRG1L sequences, and only 5 IRG1 sequences (<1.59%; Fig. S1 and Fig. 2).

To further establish the paralogous relationship between IRG1L and IRG1, we performed a manual synteny analysis to probe the genomic context surrounding pIRG1, IRG1L and IRG1 in the representative species from the major metazoan taxa that include well-studied model organisms (Fig. 2). This subset of the tree retains both the branch topography (pIRG1, IRG1L and IRG1) and the enrichment of MTS in the pIRG1 and IRG1L homologs (Fig. 2). We then manually annotated and compared the coding genes in the genomic region harboring pIRG1/IRG1L/IRG1. Although we did not detect obvious functionally related gene clustering^44^, the synteny surrounding IRG1 appears to have converged to a more highly conserved pattern than that of IRG1L, both in the number of the syntenic genes and their directionality (Fig. 2).

Four closely examined genes syntenic to pIRG1 appear to contain the same protein domains as the genes syntenic to IRG1L, but not the genes syntenic to IRG1, which we labeled based on their conserved protein domains: VOC, YrdC, 2Fe (ii) oxy and GAT (Fig. 2). The exception is the POZ domain protein that is found in the oyster synteny but are fused to BTB domain in the human gene KCTD12 that is syntenic to human IRG1. Therefore, pIRG1 might have a more similar synteny to IRG1L than to IRG1.

To experimentally study sub-mitochondrial localization, we applied protease K treatment of mitochondrial fraction of HEK293 cells expressing zebrafish IRG1L that contains a predicted MTS. Consistent with the *in silico* analysis, zebrafish IRG1L can only be digested in the presence of a strong detergent Triton X-100, similar to a matrix protein marker HSP60, confirming its matrix localization (Fig. 3A). In contrast, a recent super-resolution microscopy and biochemistry using protease treatment identified that human IRG1 lacking MTS is not localized to matrix, despite a mitochondrial enrichment^45^. We then engineered matrix-targeted human IRG1 by fusing IRG1 with a model MTS, and demonstrated its matrix localization (Fig. 3B). Together, the matrix localization of zebrafish IRG1L might suggest a broader matrix localization of pIRG1 and other IRG1Ls with MTS but not majority of IRG1 that lack MTS. This loss of MTS in the IRG1 and loss of matrix-targeted IRG1L in the modern mammals might confer a fitness advantage by redirecting IRG1 to cytoplasm and alleviating itaconate’s direct inhibition of the mitochondrial TCA cycle energy metabolism.

**Figure 3.**
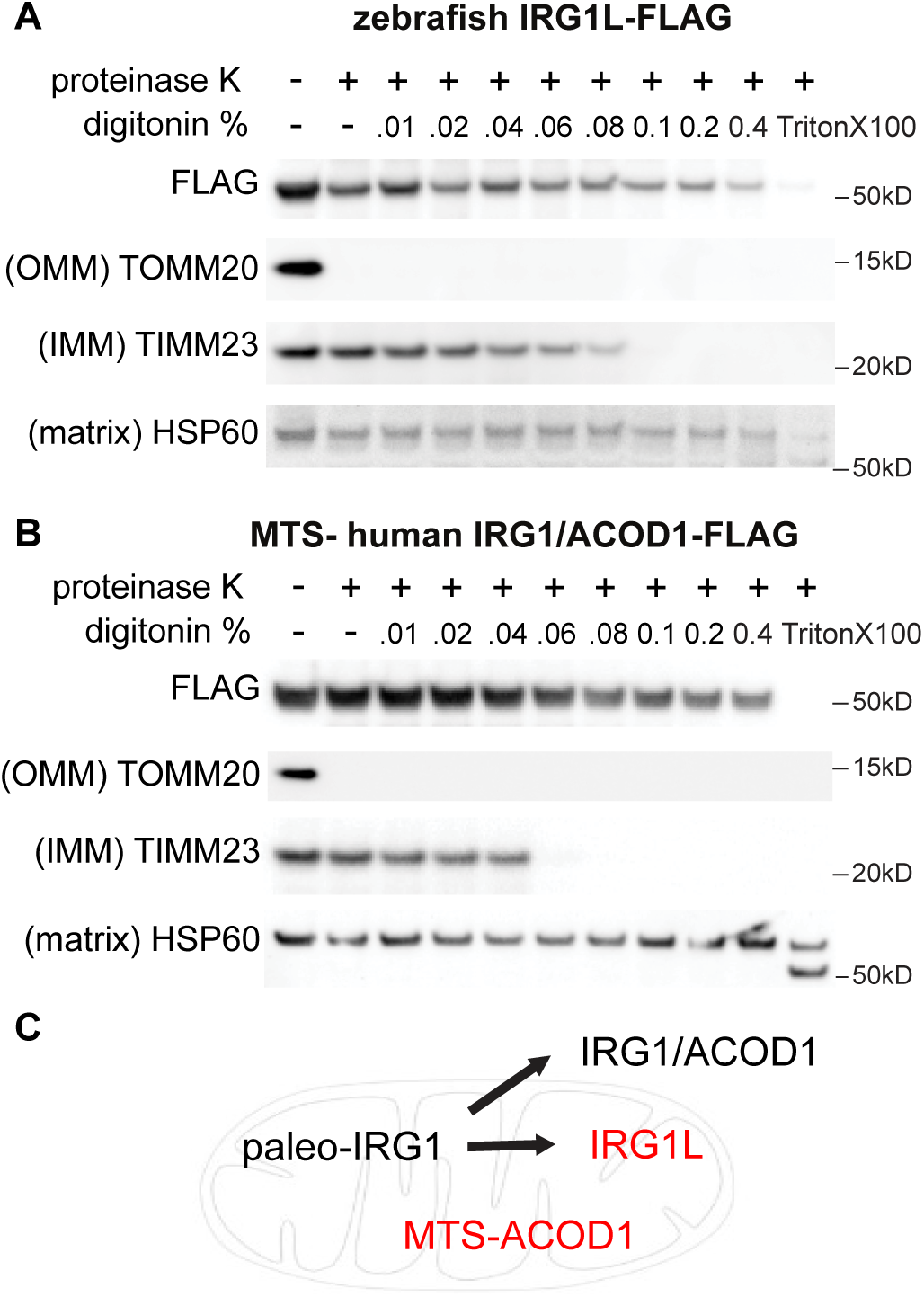
Mitochondrial matrix localization of zebrafish IRG1L. Limited proteolysis of mitochondria from HEK293T cells expressing zebrafish IRG1L (A) and human IRG1 with engineered mitochondrial targeting sequence (MTS) (B). **C**, diagram summarizing the representative submitochondrial localization of paleo-IRG1, IRG1L, IRG1/ACOD1 and engineered MTS-ACOD1.

To confirm that pIRG1 and IRG1L are indeed cis-aconitate decarboxylase and are thus implicated in innate immunity similar to human IRG1, we performed both *in vitro* enzymology to study the recombinant zebrafish IRG1L protein, as well as *ex vivo* cellular and tissue-level analysis of paleo-IRG1 in eastern oysters (*Crassostrea virginica*) and amphioxus (*Branchiostoma floridae*).

First, *in vitro* enzymology using the zebrafish IRG1L recombinant protein confirmed the same cis-aconitate decarboxylase activity that is reported for human IRG1 and *A. terreus* ACOD^26^. We purified N-terminally His-tagged recombinant zebrafish IRG1L protein (aa 32-466) lacking the predicted mitochondrial targeting sequence (Fig. 4A). Incubating this recombinant protein with aconitate *in vitro* produced itaconate in an enzyme- and time-dependent manner, detected by LC-MS (Fig. 4B and 4C).

**Figure 4.**
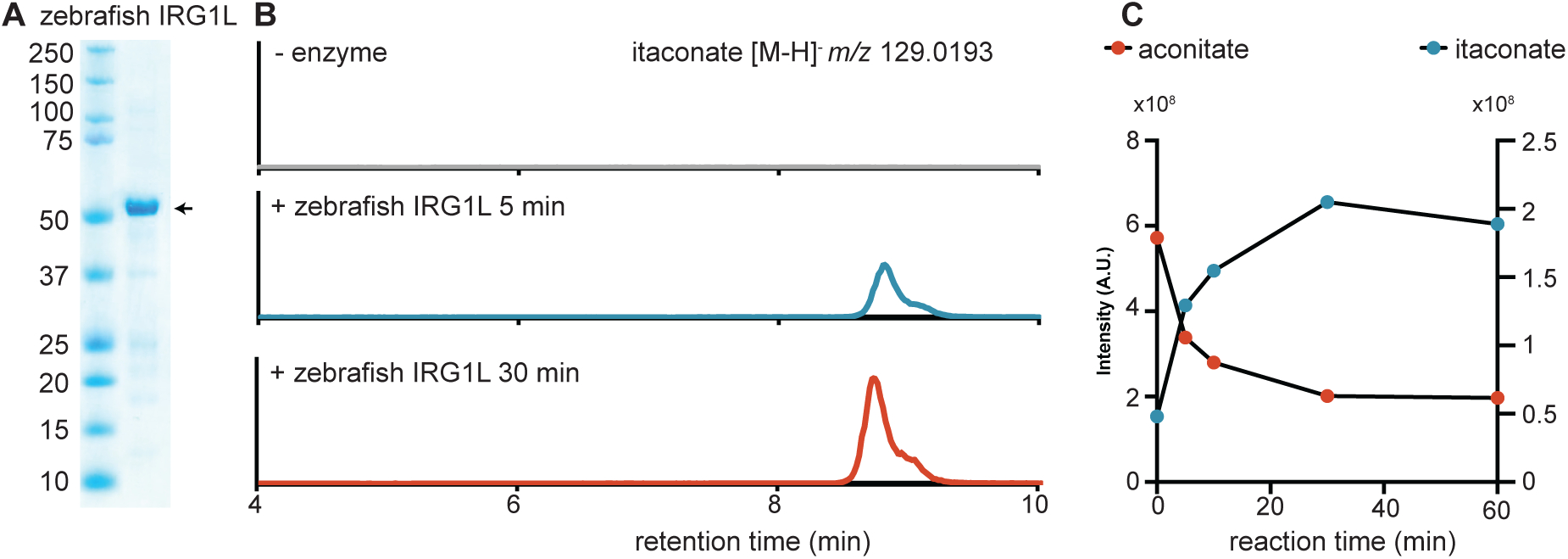
Zebrafish IRG1L is a *cis*-aconitate decarboxylase producing itaconate. (A) Recombinant His-tagged zebrafish IRG1L was expressed from *E. coli* and purified by affinity purification. (B-C) LC-MS analysis confirms itaconate production from *cis*-aconitate (1mM) by zebrafish IRG1L in a time-, enzyme-dependent manner.

To probe a potentially ancient role of pIRG1 in innate immunity, we then analyzed pIRG1 in the most distant metazoan known to contain the IRG1 gene, mollusks, specifically eastern oysters (*C. virginica*). Under resting conditions, the pIRG1 mRNA can be detected from several major oyster tissues including gills, mantle, stomach, muscles, and heart (Fig. 5A and 5B), however, a strikingly high level of pIRG1 was detected in the hemocytes (P < 0.0001 relative to other tissues; Fig. 5B), the cells in the oyster hemolymph extracted from the adductor muscles^46^. The adult oysters contain a semi-open circulatory system in which the hemolymph (analogous to blood) directly bathes other organ systems^47–49^. The oyster hemocytes exhibit dual functionality in both nutrient acquisition and innate immune defense via phagocytic properties, resembling the macrophages of the starfish larvae observed by Elie Metchnikoff^50,51^.

**Figure 5.**
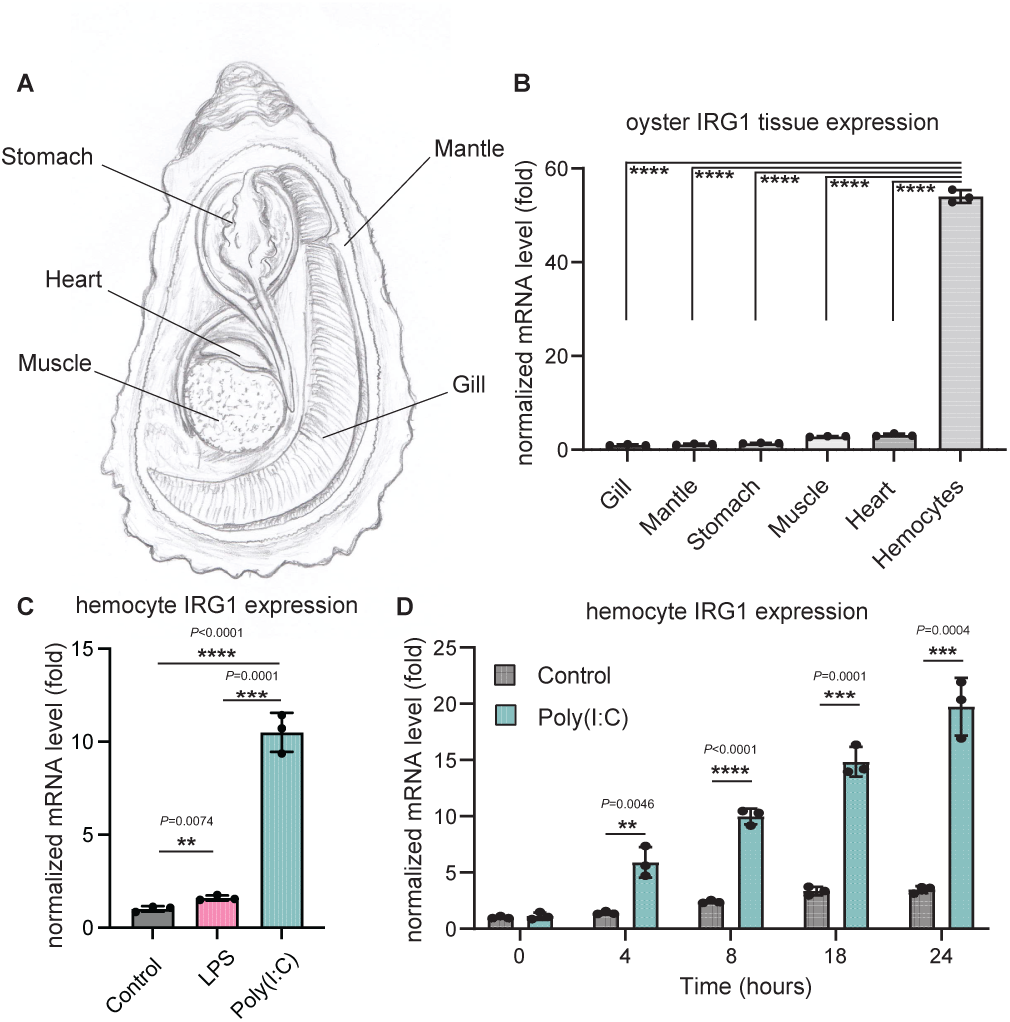
The expression of paleo-IRG1 in eastern oysters is stimulated by Poly(I:C). (A) oyster anatomy. (B) Quantitative PCR showing unstimulated oyster (*n* = 5) pIRG1 tissue expression (*P*_Hemocytes/other tissue_ < 0.0001). (C) The pIRG1 expression in the oyster hemocytes is induced Poly(I:C) at 8 hrs shown by qPCR. (D) The time course of pIRG1 expression induction in the oyster hemocytes by Poly(I:C) (10 ug/ml) stimulation. Data are shown as mean ± s.d.

To further explore pIRG1 in oyster innate immunity, we isolated and cultured pooled oyster hemocytes and measured pIRG1 mRNA level in response to a panel of innate immune inducers: heat-killed gram-negative bacterial pathogen *E. coli* (HKEB), heat-killed gram-positive bacterial pathogen *S. aureus* (HKSA), zymosan (mimicking fungal infection), and Poly(I:C) (mimicking viral infection) for 8 hours (Fig. S2). We found that oyster pIRG1 was minimally induced by LPS (mimicking gram-negative bacterial infection), but was significantly upregulated by poly(I:C) stimulation (Fig. S2 and Fig. 5E), and that Poly(I:C) induction upregulated pIRG1 expression up to 24-hr stimulation (Fig. 5D). The finding suggested an involvement of pIRG1 in oyster innate immunity.

We then investigated pIRG1 in the basal chordate amphioxus *B. floridae*. Amphioxus, or lancelets, belong to the subphylum cephalochordata, which is thought to have shared a common ancestor with more derived chordates > 500 mya^52^. This 4-7 cm long, translucent, fish-like animal contains a notochord, but not vertebrae, is considered a basal chordate and a model organism to study origins of vertebrates^53^. Lacking circulatory cells, the cellular immunity in the adult animals is provided by tissue-resident macrophage-like coelomocytes, as reported in the intestine^54^. To probe amphioxus pIRG1 in innate immunity, the adult animals were stimulated with LPS and Poly(I:C) by oral administration, and then recovered in salt water for 24 hrs prior to tissue dissection and analysis for pIRG1 transcript (Fig. 6A). Paleo-IRG1 is barely detected in the major tissues including the gills, hepatocecum, muscle and intestine in the control animals (Fig. 6B), where the inconsistent detection with very large amplification cycle numbers (>35) precluded an accurate normalization. In comparison, the pIRG1 transcript was dramatically upregulated and consistently detected in all tissues after both LPS and Poly(I:C) stimulation (Fig. 6B). A transcriptional induction of amphioxus pIRG1 by LPS and Poly(I:C) supports the role of pIRG1 and itaconate biochemistry in innate immunity early in metazoans.

**Figure 6.**
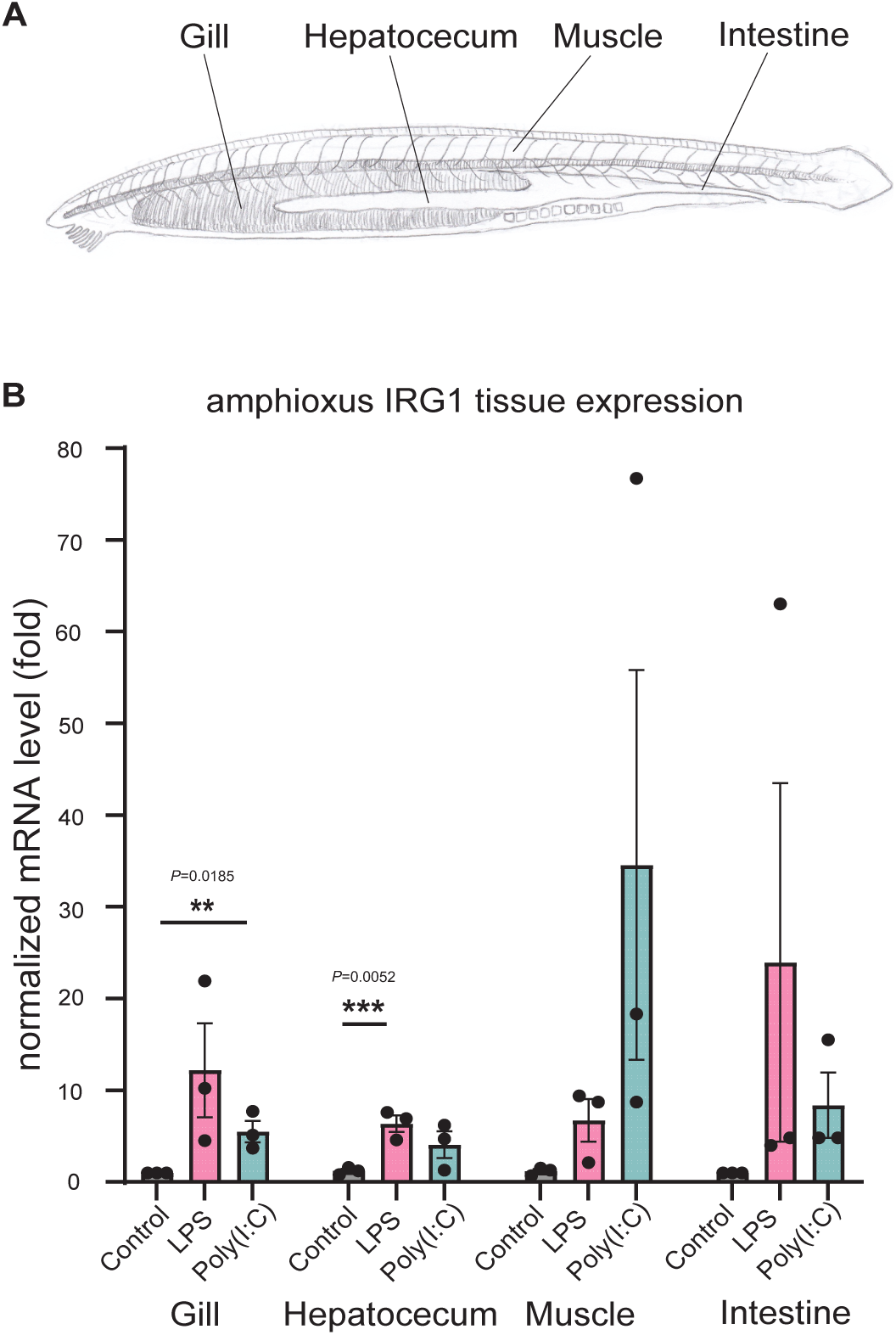
The paleo-IRG1 in amphioxus is implicated in innate immunity. (A) amphioxus anatomy. (B) Quantitative PCR from amphioxus tissues showing that pIRG1 is barely expressed under unstimulated condition (*n* =3), but can be strongly induced by both LPS (50 ug/mL; *n* =3) and Poly (I:C) (1 mg/mL; *n* = 3) upon oral injection for 24 hrs. Each data point represents tissue from one animal shown as mean ± s.d. The large deviance is likely due to the low and variable mRNA level under basal condition.

## Discussion

Our experimental studies of IRG1 in oysters and amphioxus, the most distant metazoan species known to contain IRG1, revealed itaconate biosynthesis is enriched in the innate immune cells (in oysters) and is induced upon innate immune stimulus (in oysters and amphioxus), suggesting an ancient and yet already specialized role of itaconate in innate immunity. The induction of oyster pIRG1 is consistent with the increased itaconate level reported in oysters upon marine herpesvirus infection^55^ and is also supported by the induced expression of pIRG1^56^ and increased itaconate^57,58^ in another marine bivalve (mussels; *Perna canaliculus*) upon infection with a gram-negative *Vibrio* strain. In these contexts, itaconate might play an antimicrobial function, as itaconate inhibits a pathogenic marine *Vibrio* strain growth *in vitro* and disruption of central carbon metabolism^59^.

Taxonomic distribution and evolutionary trajectory of metazoan IRG1 provide insights how molecular adaptations might regulate itaconate biochemistry for innate immunity. The frequent loss of IRG1 in metazoan taxa, particularly the loss of IRG1L paralog in mammalian taxa, clearly suggests itaconate biosynthesis is dispensable for the host’s organismal central energy metabolism. Due to potentially detrimental effects of itaconate in inhibiting host energy metabolism, itaconate production might reduce host fitness and may therefore be subject to negative selection and/or stringent regulatory strategies in evolution, a notion prompted by several observations in this study.

First, we identified that the mitochondrial targeting sequence (MTS) is frequently detected in pIRG1 and IRG1L, but is lost in IRG1, the only gene copy that is retained in the eutherian mammals. The loss of IRG1L in the therians coupled with the loss of MTS in IRG1 would relocate the catalysis outside the mitochondria and prevent direct inhibition of the mitochondrial matrix-localized energy metabolism. Despite lacking MTS, mammalian IRG1 remains associated with the mitochondria with unknown mechanism^59–61^. A recent identification of a direct IRG1-mediated mitochondria-vacuole interface highlighted the importance of this efficient mitochondria-derived itaconate production in host defense against intracellular pathogen *Salmonella*^45^. Second, despite a conserved role of itaconate in metazoan innate immunity, IRG1 transcription in mammals is more tightly regulated than that of the basal metazoans. While the mammalian IRG1 is almost exclusively restricted to LPS-activated TLR4 and interferon signaling in macrophages^60,62^, the pIRG1s in oysters and in amphioxus appear to be transcriptionally regulated with less stringency, including a basally expressed pIRG1 in oysters’ tissues and hemocytes without any stimulation.

We reported a dramatic transcriptional induction of oyster pIRG1 by poly(I:C) and the low level of induction by LPS stimulation, as well as a broader, dual induction of amphioxus pIRG1 by both poly(I:C) and LPS stimulation. Our report on poly(I:C)-induced oyster pIRG1 expression is supported by the transcriptomics analysis in mussel hemocytes upon screening for a panel of pathogen-associated molecular patterns (PAMPs), where mussel pIRG1 induction is restricted to poly(I:C) stimulation^63^. Lophotrochozoa (including marine bivalves) and amphioxus contain rudimentary TLR sequences^64–67^, however, it remains unknown if these rudimentary TLRs function as pattern recognition receptors (PRRs) with the same degree of specificity and downstream signaling as their mammalian counterparts^67,68^. Notably, the first identified Drosophila Toll-receptor does not recognize foreign PAMPs despite the involvement in host resistance against fungal and bacterial infection^69^. The strikingly high upregulation of pIRG1 from Poly(I:C) stimulation relative to LPS leads us to speculate that pIRG1 in oysters and amphioxus might be induced by a non-TLR, membrane-bound PRRs. For instance, zebrafish IRG1 is regulated by cooperation between glucocorticoid receptors and the JAK/STAT pathway^20^. Future multispecies analysis on the immune signaling pathways and transcriptional factors that induce IRG1 expression in non-traditional model organisms is vital to answer these questions.

Investigation of three metazoan subbranches, pIRG1, IRG1L and IRG1, support the notion that genome duplication, horizontal gene transfer and gene loss provide powerful evolutionary forces to innovate gene function and regulation^70^. Our analysis is consistent with the notion that whole genome duplication (WGD) in the vertebrate lineage that produced the paralogous IRG1L and IRG1, in which IRG1L is subsequently lost in viviparous mammals. The timings of the two rounds (2R) WGD of the early vertebrate lineage have been reported before^71^ and after^41^ the gnathostome-agnathan divergence occurring around 465-428 Ma. Our findings that lamprey has two pIRG1s without either IRG1 and IRG1L sequence would agree with 2R WGD and the one after the divergence of agnathans and gnathostomes gave rise to IRG1 and IRG1L. The relative similarity of IRG1L and pIRG1 synteny and the presence of MTS would support the notion that the genome duplication and loss of IRG1L enable a tightened regulation of itaconate biosynthesis exerted by extra-mitochondrial IRG1 that is subsequently retained in eutherian mammals.

Our exploration of the molecular evolution of itaconate biosynthesis presents another case of horizontal gene transfer in eukaryotes from prokaryotic source in the context of innate immunity. The potential independent acquisition of the metazoan IRG1 and the fungi *A. terreus* ACOD suggests at least one, and perhaps two horizontal gene transfer (HGT) events from bacteria to eukaryotes in the evolutionary history of IRG1. The exclusive presence of ACOD in *Aspergillus* taxa within fungi that are nested within other prokaryotic sequences would strongly suggest a HGT in *Aspergillus*. Our studies of the metazoan IRG1 can be explained by two scenarios: first, the metazoan IRG1 is derived from the last common ancestor of eukaryotes and subsequently lost several times in major eukaryotic taxa including plants, most fungi, and ecdysozoa; and second, the metazoan ancestor’s IRG1 is horizontally transferred directly from bacteria. The observation that the closest branch next to metazoan IRG1 (Fig. 1C) contains sequences from alphaproteobacteria might support the former. However, the frequent local gene duplication in pIRG1 (e.g., in oysters and in sea lampreys) and IRG1L (in xenopus) suggests pIRG1/IRG1L genomic region might be highly mobile, indicative of a more recent HGT event — an analysis of mobile gene elements near pIRG1 would help resolve this question. Notably, a previous systems-level analysis of eukaryotic orthologous groups (KOGs) also highlighted CAD sequences as one of the unique eukaryotic sequences with unexpected phyletic patterns, supporting putative independent HGT events^72^. In microorganisms, the study on itaconate production in immunometabolism of other bacterial and fungi species might also indicate a universal role of itaconate in innate immunity.

## Methods

### Phylogenetic analysis of the IRG1 orthologs

Human IRG1 was manually queried against the NCBI database using BLASTp and tBLASTn^73^ across the tree of life and were subsampled by taxonomic order during May-July 2020.

Sequences from metazoan taxa were restricted to Expectation values below 1e-10. To maximize sequence diversity in more distantly related sequences from fungi, archaea, and prokaryotes inclusion criteria was relaxed to 0.001. In total, 2,297 sequences were identified. Duplicate sequences were identified and removed. A full phylogeny of all sequences was constructed using MAFFT (v7.471)^74^. IQTree (v2.1.2)^75^ and ModelFinder^76^ were used to find the best fit substitution model and branch support values were obtained by ultrafast bootstrapping^77^. Figtree (v1.4.4) and iTOL (v6.5.2)^78^ were used for tree visualization.

### Analysis of dubious sequences

Initial sequence surveys revealed only nine orthologs in Arthropoda, all in *Drosophila* genera, three orthologs in Plantae: one in *Ziziphus* and two in *Quercus* generas. Reciprocal BLASTp and phylogenetic analysis indicated likely sequence contamination by *Acetobacter aceti* in Drosophila sequences, a black yeast-like fungal pathogen in the *Quercus* sequence, and *Belnapia* contamination in the *Ziziphus* sequences. Therefore, these sequences were removed.

### Identification of mitochondrial targeting sequences (MTS)

All the metazoan IRG1 sequences were analyzed using Target-P (v2.0)^79^ to predict the presence of N-terminal MTS. Default settings were used, and a likelihood >0.5 was considered a predicted MTS.

### Synteny analysis

Representative genes from the model organisms from each major metazoan taxa possessing an IRG1 ortholog were manually investigated for syntenic genes using NCBI Genome Data Viewer^80^. The five nearest annotated genes both upstream and downstream were analyzed using Pfam^81^ and NCBI Domain Architecture Retrieval Tool^82^ to predict domain architecture. In instances where local duplication is present, more than the five nearest genes were recorded as there were multiple nearby IRG1 homologs (e.g., *Xenopus laevis*).

### Plasmids and cloning

A N-terminal His-tagged *Danio rerio* IRG1L sequence lacking the N-terminus predicted mitochondrial targeting sequence (Δ31aa) was codon-optimized for bacterial expression, custom synthesized and subcloned into pET-28a(+)-TEV vector using Nde1 and BamH1 sites (GenScript).

### His-Zebrafish IRG1L (aa 32-466)

MGSSHHHHHHSSGENLYFQGHM

APEETVTSSFGRFIQSVQPKNLTPTVLQRSKRMVLDSIGVGLVGSTTEVFDLALQHCQHMYASDDISFVYGRQGVKLSPTLAAFVNGVAAHSMDFDDTWHPATHPSGAVLPALLAISDMLHSNSKPSGLDFLTAFNVGIEIQGRLMRFSNEAHNIPKRFHPPSVVGTLGSAAACARLLSLDRNQASNALAIAASLAGAPMANAATQSKPLHIGNASRLGLEAALLASRGLEASPLVLDAVPGVAGFSAFYEDYAPRPIGSPEEDEHSFLIESQDIAFKRFPAHLGMHWIADAASVVHKTLVGLKGGSISPNLVQDILLRVPLSKYINRPFPESEHQARHSFQFNACTALLDGEVTVQSFHPEAIQRPELHALLSRVRLEHPSDNPANFNIMYGEVEVTLVTGDVLRGRCDTFYGHWRNPLSQESLHKKFRNNAGTVLSTEKVERLIEAVESLDRSDDCKQLLAQLQ*

gcgccggaggaaaccgtgaccagcagcttcggtcgttttatccagagcgttcaaccgaagaacctgaccccgaccgtgctgcagcgtagc aaacgtatggttctggacagcattggtgtgggcctggttggcagcaccaccgaggtgttcgatctggcgctgcagcactgccaacacatgta cgcgagcgacgatatcagctttgtgtatggtcgtcaaggcgttaagctgagcccgaccctggcggcgtttgtgaacggtgttgcggcgcac agcatggactttgatgatacctggcatccggcgacccacccgagcggtgcggtgctgccggcgctgctggcgatcagcgacatgctgcac agcaacagcaagccgagcggtctggatttcctgaccgcgtttaacgttggtatcgagattcagggccgtctgatgcgtttcagcaacgaagc gcacaacattccgaaacgttttcacccgccgagcgtggttggtaccctgggcagcgcggcggcgtgcgcgcgtctgctgagcctggatcg taaccaggcgagcaacgcgctggcgattgcggcgagcctggcgggtgcgccgatggcgaacgcggcgacccaaagcaaaccgctgc acattggtaacgcgagccgtctgggtctggaggcggcgctgctggcgagccgtggtctggaagcgagcccgctggtgctggatgcggtg ccgggtgttgcgggtttcagcgcgttttacgaggattatgcgccgcgtccgattggcagcccggaggaagacgagcacagcttcctgatcg aaagccaggatattgcgttcaagcgttttccggcgcacctgggtatgcactggattgcggacgcggcgagcgtggttcacaagaccctggt tggcctgaaaggtggcagcatcagcccgaacctggtgcaagatattctgctgcgtgttccgctgagcaaatacatcaaccgtccgttcccgg agagcgaacaccaggcgcgtcacagcttccaatttaacgcgtgcaccgcgctgctggatggtgaggtgaccgttcagagctttcacccgg aggcgatccaacgtccggaactgcacgcgctgctgagccgtgtgcgtctggaacacccgagcgataacccggcgaacttcaacattatgt acggcgaggtggaagttaccctggtgaccggtgacgttctgcgtggccgttgcgataccttctatggtcactggcgtaacccgctgagcca ggaaagcctgcacaagaaatttcgtaacaacgcgggcaccgtgctgagcaccgagaaggttgaacgtctgattgaggcggttgaaagcct ggaccgtagcgacgattgcaaacaactgctggcgcagctgcaataa

For mitochondrial localization assay in the HEK293 cells, sequences were codon-optimized for mammalian expression, custom synthesized and subcloned into in lentiviral pLys1 vector (Addgene #50058) using Nhe1/Xma1or EcoRV/Xma1 (for MTS-human IRG1) sites (GenScript).

### Human IRG1/ACOD1-FLAG

MMLKSITESFATAIHGLKVGHLTDRVIQRSKRMILDTLGAGFLGTTTEVFHIASQYSKIYSSNISSTVWGQPDIRLPPTYAAFVNGVAIHSMDFDDTWHPATHPSGAVLPVLTALAEALPRSPKFSGLDLLLAFNVGIEVQGRLLHFAKEANDMPKRFHPPSVVGTLGSAAAASKFLGLSSTKCREALAIAVSHAGAPMANAATQTKPLHIGNAAKHGIEAAFLAMLGLQGNKQVLD LEAGFGAFYANYSPKVLPSIASYSWLLDQQDVAFKRFPAHLSTHWVADAAASVRKHLVAERALLPTDYIKRIVLRIPNVQYVNRPFPVSEHEARHSFQYVACAMLLDGGITVPSFHECQINRPQVRELLSKVELEYPPDNLPSFNILYCEISVTLKDGATFTDRSDTFYGHWRKPLSQEDLEEKFRANASKMLSWDTVESLIKIVKNLEDLEDCSVLTTLLKGPSPPEVASNSPACNNS ITNLS PGGGSGGSGGSMDYKDDDDK*

gctagcgccaccatgatgctgaaaagcatcacagagagcttcgcgaccgccattcacggactgaaggtgggccacctgacagatagagtgatccaaagatctaaaaggatgatcctggatacactgggcgctggattcctgggtaccaccaccgaggtgttccatatcgccagccagtacagcaagatctacagcagcaacatctctagcacagtgtggggacagcctgacatcagactgccacctacctacgccgccttcgtgaacggcgtggccatccacagcatggacttcgacgacacatggcaccccgccacccacccttccggagccgtgctgcctgtgttaacagccctggctgaagccctgccaagatcccctaagttcagcggcctggacctgctgctggccttcaacgtgggcatcgaggtgcaaggcagactgctgcatttcgccaaggaagccaacgacatgcctaagcggttccaccctcctagcgtggtgggcacactgggcagcgccgcggccgcgtccaagtttctgggcctgagcagcaccaagtgcagagaggctctggccatcgccgtgtctcacgccggcgcccctatggccaatgccgccacgcagaccaagcccctgcacatcggcaatgccgccaagcacggcatcgaggccgcctttctggccatgctgggcctgcagggcaacaagcaggtgctcgacctggaagccggcttcggcgccttctacgccaattacagcccaaaagtgctgcctagcatcgcctcatattcttggctgctggatcagcaggacgtggccttcaagcggtttccagcccacctgagcacccactgggtggctgatgccgctgccagcgtcagaaagcacctggtggcagaaagagccctgctgcccaccgactacatcaaacggattgtgctgagaatacccaacgtgcagtacgtgaaccggcccttccctgtcagcgagcacgaggctagacactcttttcagtatgtggcttgtgccatgctactggacggcggaatcaccgtgccttcttttcacgagtgccagatcaacagacctcaggtccgcgaactgctgtccaaggtggaactcgaatacccccccgataacctgcctagcttcaacatcctgtactgcgagatcagcgtgaccctgaaggacggcgccacattcaccgacagaagcgacaccttctacggccactggcggaagcctctgtctcaggaggatctggaggaaaagttcagagctaacgcctccaaaatgctgagctgggacaccgtggaaagcctgatcaagatcgttaagaacctggaggacctg gaggactgcagcgtgttgacaaccctgctgaagggcccttcccctcccgaggttgctagtaatagccctgcctgtaacaacagcatcaccaacctgtcccccgggggtggatctggtggatctggtggatctatggattacaaggatgacgatgacaagtaa

### MTS-human IRG1L-FLAG (with MTS from cytochrome C oxidase subunit 4I1 (COX4I1))

MLATRVFSLVGKRAISTSVCVRAHAS

MMLKSITESFATAIHGLKVGHLTDRVIQRSKRMILDTLGAGFLGTTTEVFHIASQYSKIYSSNISSTVWGQPDIRLPPTYAAFVNGVAIHSMDFDDTWHPATHPSGAVLPVLTALAEALPRSPKFSGLDLLLAFNVGIEVQGRLLHFAKEANDMPKRFHPPSVVGTLGSAAAASKFLGLSSTKCREALAIAVSHAGAPMANAATQTKPLHIGNAAKHGIEAAFLAMLGLQGNKQVLDLEAGFGAFYANYSPKVLPSIASYSWLLDQQDVAFKRFPAHLSTHWVADAAASVRKHLVAERALLPTDYIKRIVLRIPNVQYVNRPFPVSEHEARHSFQYVACAMLLDGGITVPSFHECQINRPQVRELLSKVELEYPPDNLPSFNILYCEISVTLKDGATFTDRSDTFYGHWRKPLSQEDLEEKFRANASKMLSWDTVESLIKIVKNLEDLEDCSVLTTLLKGPSPPEVASNSPACNNS ITNLS PGGGSGGSGGSMDYKDDDDK*

gatatcgccaccatgctggctacaagagtgttcagcctggtcggcaagagagccatcagcacctctgtgtgcgtgcgggcccacgctagcatgatgctgaaaagcatcacagagagcttcgcgaccgccattcacggactgaaggtgggccacctgacagatagagtgatccaaagatctaaaaggatgatcctggatacactgggcgctggattcctgggtaccaccaccgaggtgttccatatcgccagccagtacagcaagatctacagcagcaacatctctagcacagtgtggggacagcctgacatcagactgccacctacctacgccgccttcgtgaacggcgtggccatccacagcatggacttcgacgacacatggcaccccgccacccacccttccggagccgtgctgcctgtgttaacagccctggctgaagccctgccaagatcccctaagttcagcggcctggacctgctgctggccttcaacgtgggcatcgaggtgcaaggcagactgctgcatttcgccaaggaagccaacgacatgcctaagcggttccaccctcctagcgtggtgggcacactgggcagcgccgcggccgcgtccaagtttctgggcctgagca gcaccaagtgcagagaggctctggccatcgccgtgtctcacgccggcgcccctatggccaatgccgccacgcagaccaagcccctgcacatcggcaatgccgccaagcacggcatcgaggccgcctttctggccatgctgggcctgcagggcaacaagcaggtgctcgacctggaagccggcttcggcgccttctacgccaattacagcccaaaagtgctgcctagcatcgcctcatattcttggctgctggatcagcaggacgtggccttcaagcggtttccagcccacctgagcacccactgggtggctgatgccgctgccagcgtcagaaagcacctggtggcagaaagagccctgctgcccaccgactacatcaaacggattgtgctgagaatacccaacgtgcagtacgtgaaccggcccttccctgtcagcgagcacgaggctagacactcttttcagtatgtggcttgtgccatgctactggacggcggaatcaccgtgccttcttttcacgagtgccagatcaacagacctcaggtccgcgaactgctgtccaaggtggaactcgaatacccccccgataacctgcctagcttcaacatcctgtactgcgagatcagcgtgaccctgaaggacggcgccacattcaccgacagaagcgacaccttctacggccactggcggaagcctctgtctcaggaggatctggaggaaaagttcagagctaacgcctccaaaatgctgagctgggacaccgtggaaagcctgatcaagatcgttaagaacctggaggacctggaggactgcagcgtgttgacaaccctgctgaagggcccttcccctcccgaggttgctagtaatagccctgcctgtaacaacagcatcaccaacctgtcccccg gg

### zebrafish IRG1L-FLAG

MLSAVQRSSRYLTSFSAARGLHKSALDVAERPAPEETVTSSFGRFIQSVQPKNLTPTVLQRSKRMVLDSIGVGLVGSTTEVFDLALQHCQHMYASDDISFVYGRQGVKLSPTLAAFVNGVAAHSMDFDDTWHPATHPSGAVLPALLAISDMLHSNSKPSGLDFLTAFNVGIEIQGRLMRFSNEAHNIPKRFHPPSVVGTLGSAAACARLLSLDRNQASNALAIAASLAGAPMANAATQSKPLHIGNASRLGLEAALLASRGLEASPLVLDAVPGVAGFSAFYEDYAPRPIGSPEEDEHSFLIESQDIAFKRFPAHLGMHWIADAASVVHKTLVGLKGGSISPNLVQDILLRVPLSKYINRPFPESEHQARHSFQFNACTALLDGEVTVQSFHPEAIQRPELHALLSRVRLEHPSDNPANFNIMYGEVEVTLVTGDVLRGRCDTFYGHWRNPLSQESLHKKFRNNAGTVLSTEKVERLIEAVESLDRSDDCKQLLAQLQ PGGGSGGSGGSMDYKDDDDK*

gctagcgccaccatgctgtctgccgttcagaggtccagcagatacctgaccagcttcagcgccgctagaggcctccacaaaagcgccctggacgtggccgagaggcctgctcctgaagagacagtgaccagcagcttcggcagattcatccagtctgtgcaacctaagaacctgacacctaccgtgctgcaacgctccaaaagaatggtgctcgactccatcggcgtgggcctggtgggcagcacaaccgaggtgttcgacttggccctgcagcactgccagcatatgtatgcctccgacgacatcagctttgtgtacggccggcagggcgtgaagctgagcccaacactggctgctttcgtgaacggcgtggccgctcatagcatggacttcgacgatacctggcaccctgccacccacccttcaggcgccgtgctccctgctctgctggctatcagcgacatgctgcattctaatagcaagccaagcgggctggacttcctgaccgcctttaacgtgggcatcgaaatccagggaagactgatgagatttagcaacgaggcccacaacatccccaagagattccacccacctagcgtggtgggcaccctgggctctgctgcagcttgtgccagactgctgagcctggacagaaaccaggccagcaacgccctggccattgccgccagtctggccggcgcccctatggccaacgccgccacacagagtaagcctctgcacatcggcaacgccagccggctgggcctggaagccgccctgctggccagcagaggcctggaagccagccctctggtcctggatgccgtgcccggcgtcgccggatttagcgccttctacgaggactacgcccctagacctatcggatctcctgaggaagatgagcacagcttcctgatcgagagccaggacatcgccttcaagcggttccccgcccacctgggcatgcactggatcgccgacgccgccagcgtggtgcacaagaccctggtgggcctgaagggcggatctatcagccccaatctggtgcaggacatcctgctgcgggtgcctctgtccaagtacatcaacagacccttccctgagagcgagcaccaggcccggcacagcttccagttcaacgcctgcaccgccctgctggatggagaggtgaccgtgcagagctttcaccccgaagcaattcaaagacccgagctgcacgccctgttgtccagagtgcggctggaacacccatctgacaaccccgctaatttcaacatcatgtacggcgaggtcgaggtgaccctggtgacaggcgacgtgctgagaggacggtgcgacaccttctacggccactggcggaatcctctgtctcaggagagcctacacaaaaagttcagaaacaacgctggtacagttctgagcaccgagaaggtggaaagactgatcgaagccgtggaaagcctggatagatctgatgattgcaagcagctgctggcccagctgcagcccggg

### Zebrafish IRG1/ACOD1-FLAG

MIRKSVTDSFGAAVSCLSTSHLTDEVIRRSKRMILDTLGVGLIGTRTPVFNTVLQCSQRQQALENSKVWGRPGSSLPPQYAAFVNGVAVHSMDFDDTWHPATHPSGAVLPALLALAETLPVKPSGLELLLAFNVGIEIQGRLLRFSKEAHNIPKRFHPPAVVGVMGSAAATAKLLRLPAAQSIAALAIACSSAGAPMANAATQTKPLHMGNAARGGLEASQLALFGLVGNTHILDL PSGFGSFYPDYVPHPLAEVTPNSHYRWVLEEQDIAQKRFPAHLGMHWVADAAIEARAKFLDKHPNADLSQIKKITLRVPSSRYVDCPLPVTEHQARHSFQFNCCTALLDGEVNVQSFSRAQMNRSVLKELLLKVELENPQDNHSSFQKMYCEITVLSTQGEMFTARCDTFYGHWRKPLSQEDLLKKFRLNASSVLPNEVLEEIIYAVDHLDTNQDCSKLWSYMHFNKHMVQRDE RRCFALA PGGGSGGSGGSMDYKDDDDK

gctagcgccaccatgatccggaaaagcgtgacagatagctttggagccgccgtgtcctgcctgtctacaagccacctgaccgatgaggtgatcagaagaagcaagcggatgatcctggacacactgggcgtgggcctgattggaacaagaacccctgtgttcaacaccgtgctgcagtgcagccaaagacagcaggccctggaaaacagcaaggtgtggggcagacccggctcttccctgcctcctcaatacgccgccttcgtgaacggcgtggccgtgcacagcatggactttgacgacacctggcaccccgccacacatcctagcggtgccgtgctgcctgctctgctggccctggctgaaaccctgccagtgaagccttctggcctggagctgctgctggcctttaatgtcggcatcgagattcagggcagactgctgagattcagcaaagaggcccacaacatccccaagagattccacccacctgctgttgtgggagtcatgggcagcgctgccgccacagccaagctgctgcggctgcctgccgcccagagcatcgccgctctggctatcgcatgcagcagcgccggagcccctatggccaacgccgccacccagaccaagcccctgcacatgggcaacgccgctagaggcggcctggaggccagccagctggctctgttcggcctggtggggaatacccacatcctggatctgccatctggcttcggctccttttaccccgactacgtgcctcaccccctggcagaagtgacccctaacagccattatagatgggtgcttgaagagcaggacatcgctcagaagcgcttccctgcccacctgggaatgcactgggtcgccgacgccgccatcgaggccagagctaagttcctggacaagcaccctaatgccgatctgtcgcagatcaagaaaatcacactgagagtgcccagctctagatacgtggactgtcctctgccagtgacagagcaccaggccagacactcttttcagttcaattgctgcaccgccctgctggacggcgaagtgaacgtgcaaagcttcagcagagcccagatgaaccggagcgtgctgaaggaactgctgctcaaagtggaactggaaaacccccaggacaaccacagctccttccagaaaatgtactgcgagatcaccgtgctgagcacccagggcgagatgttcaccgccaggtgtgataccttctacggccactggcggaagcctctgtctcaggaggacctgcttaagaagttccggctgaacgccagcagcgtgctccctaacgaggtcctggaagagatcatctacgccgtggaccacctggata ccaaccaggattgttccaagctgtggtcctacatgcatttcaacaagcacatggtgcagcgggacgagcggagatgcttcgccctggcccc cggg

### Submitochondrial localization by limited proteolysis

Raw mitochondrial fraction from HEK293 cells stably expressing zebrafish IRG1L-FLAG or MTS-human IRG1/ACOD1-FLAG were subjected to limited proteolysis following the previous protocol^83^. Specifically, 30 million cells were washed with cold PBS, scraped into 1 ml ice cold mito isolation buffer (10 mM Tris-HCl, 1 mM EGTA-Tris, 200 mM sucrose, pH 7.4) that also contains sub-optimal concentration of protease inhibitor (cOmplete™, Mini, EDTA-free Protease Inhibitor Cocktail, Roche) at 1 mini tablet in 30 ml isolation buffer, and homogenized with 20 strokes in 2 ml douncer homogenizer (KIMBLE). The homogenates were spun at 600 g for 10 min at 4 °C, the supernatants were spun at 7000 g for 10 min at 4 °C, and the mitos in the pellets were resuspended in 250 μl mito isolation buffer with no protease inhibitor. The mito homogenates were spun again at 600 g for 3 min at 4 °C to remove nucleus, and aliquoted into PCR tubes at approximately 20 μg protein concentration. The mitos were treated with 100 μg/ml proteinase K (Millipore Sigma, P2308) in the presence of increasing concentrations of digitonin (Millipore Sigma, BN2006) at 0.01, 0.02, 0.04, 0.06, 0.08, 0.1, 0.2 and 0.4% digitonin, or 1% Triton X-100 for 15 min at room temperature. 7 mM PMSF was added to inactivate proteinase K for 5 min prior to SDS-PAGE analysis and western blotting.

### Zebrafish IRG1L protein production

The His-Zebrafish IRG1L plasmid was transformed in BL21 Star™ (DE3) Chemically Competent E. coli (Invitrogen c6010-03), grown at 225 rpm at 37 ℃ to OD600 0.6. Cells were then induced with 0.1 mM β-D-1-thiogalactopyranoside (IPTG) for 6 hours at 18 ℃. Cells were lysed by sonication (Qsonica Q500 with microtip) in 200 mL lysis buffer (20mM HEPES pH 7.4, 500 mM NaCl, 10% glycerol v/v, 1 mM TCEP (pH 7.4), 100 U/mL DNase, 0.1 mg/mL Lysozyme, and cOmplete mini protease inhibitor cocktail) using 2 sec on and 2 sec off pulses at 25% power for 2 minutes. The recombinant IRG1-like protein was enriched using cobalt TALON Metal Affinity Resin (Takara Bio #635501) with gentle shaking for 1 hour at 4 ℃ and washed twice by centrifugation for 3 min at 3500 rpm at 4 ℃ and resuspension in the washing buffer (20 mM Tris pH 8.0, 150 mM NaCl, 20 mM Imidazole, 1 mM TCEP). Protein was eluted from the affinity resin in the elution buffer (20 mM Tris pH 8.0, 150 mM NaCl, 500 mM Imidazole, 1 mM TCEP, 10% glycerol). The recombinant protein was then concentrated using centrifugal filter units (Amicon Ultra-4, Millipore) and dialyzed using Slide-A-Lyzer dialysis cassette into storage buffer (20 mM Tris pH 8.0, 150 mM NaCl, 20% glycerol, 1 mM TCEP) overnight.

### Enzymology

The enzyme assay for the zebrafish IRG1L was performed by incubating 7 µg recombinant protein with 1 mM cis-aconitate (Sigma A3412-1G) in 300 µl reaction buffer (10 mM HEPES, 150 mM NaCl, 0.1 mM TCEP, 10% glycerol, pH 6.5) at 37 ℃. Aliquots of 50 µl were taken at 0, 10, 30, and 60 minutes and were immediately quenched with 2 % formic acid, vortexed, and centrifuged at 14,000 rpm for 20 minutes at 4 ℃. The supernatant was then transferred to a new centrifuge tube and mixed 1:9 with a 3:1 acetonitrile:methanol solvent for LC-MS analysis following the established polar metabolite profiling protocol in the lab^84^. Itaconate production was confirmed using chemical standard itaconic acid (Sigma Aldrich #I29204).

### Mollusk pIRG1 expression

Eastern oysters (*Crassostrea virginica*, Woods Hole Marine Biological Laboratory) were acclimated in filtered sea water overnight after arrival. To determine unstimulated basal pIRG1 expression, oysters (*n* = 5) were dissected on ice and 50 mg wet tissue mass from the gill, mantle, stomach, adductor muscle, and heart were collected. Tissue was washed in PBS, then homogenized in RL+BME buffer (Qiagen). mRNA extraction was performed per manufacturer’s instruction using the RNeasy mini kit (Qiagen). The mRNA concentration was determined using a Nanodrop and cDNA libraries were prepared using SuperScriptIII First-Strand Synthesis Super Mix for qRT-PCR (Thermo Fisher Scientific).

### Mollusk genes quantitative PCR

The control gene EF1a for qPCR was identified from the literature^85^. Primers for EF1a and pIRG1 were designed using primer-BLAST (NCBI) and synthesized from Integrated DNA technologies. The pilot qPCR and Sanger sequencing experiments were conducted to validate the primers for pIRG1 and EF1a. qPCR was performed using PowerUp SYBR Green Master Mix (Thermo Fisher A25742) in a 96-well format with Thermo Fisher QuantStudio 6Flex instrument.

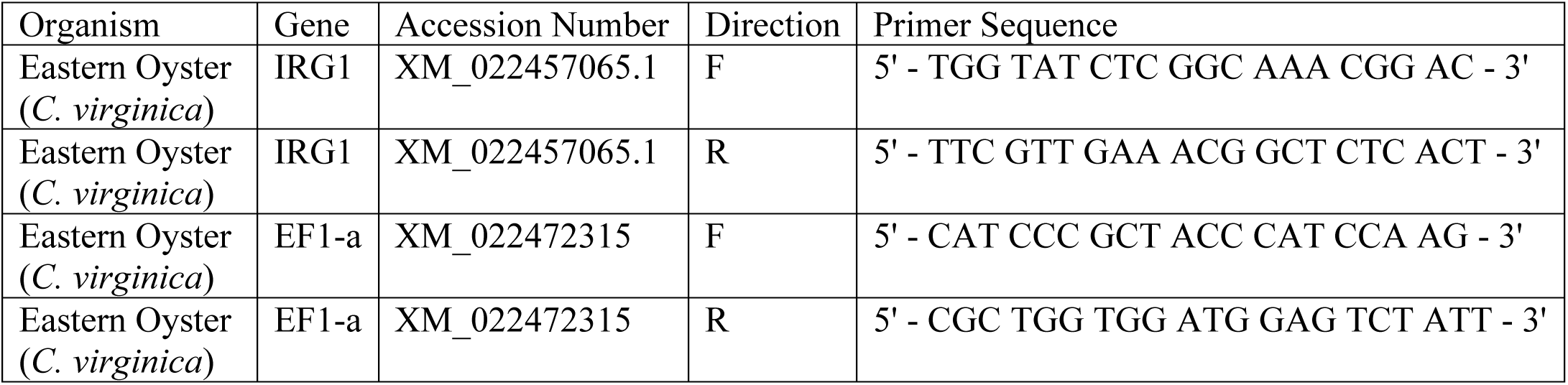

### Mollusk Hemocyte Assays

For all hemocyte assays, live eastern oysters were notched and hemolymph extracted by syringe from the adductor muscles following published protocol^46^. Hemolymph from 10 oysters was pooled on ice, diluted 1:2 with decontamination washing buffer (PBS, 100 µg/mL penicillin-streptomycin), pipetted into Corning Falcon 60 mm tissue culture plates (Thermo Fisher) and were incubated in the dark at room temperature for 30 min. Cells were then inspected under the microscope to ensure adhesion, and decontamination buffer was aspirated and replaced with 5 mL culture media (1:1 OptiMEM media: filtered sea water, 50 µg/mL penicillin-streptomycin).

The cultured hemocytes were treated with zymosan (100 µg/mL, Invivogen tlrl-zyd), heat-killed *E. coli* (1x10^7^ PFU, Invivogen tlrl-hkeb2), heat-killed *S. aureus* (1x10^7^ PFU, Invivogen tlrl- hksa), and Poly(I:C) (10 µg/mL final concentration, Invivogen tlrl-pic), and filtered sea water (control) for 8 hrs in the dark at room temperature approximately 20 ℃, or Poly(I:C) and LPS E. coli 0111:B4 (1 µg/mL final concentration, Sigma LPS25) for indicated time. After treatments, cells were directly lysed and analyzed as described above.

### Amphioxus pIRG1 expression

Amphioxus (*Branchiostoma floridae*, Gulf Specimen Marine Lab) were kept in the filtered sea water in the dark at room temperature. Three amphioxus each condition were treated by oral injection with LPS E. coli 0111:B4 (50 µg/mL final concentration, Sigma LPS25), Poly(I:C)(1 mg/mL, Invivogen tlrl-pic), or control (filtered seawater). Specifically, amphioxus were cold-immobilized and a 25 gauge needle was used to carefully administer 100 µl stimulus solution through the oral hood. Individuals were immediately placed back into their containers at room temperature and recovered overnight. Individuals were netted, cold immobilized and dissected 24 hr post treatment. Gill, muscle, hepatocecum and intestine tissue were collected and washed in PBS before tissue lysis and mRNA expression analysis.

Candidate reference genes were identified from literature^86^. Primers for pIRG1 and reference genes S20 and EF1a were designed using NCBI primer-BLAST, and synthesized (Integrated DNA technologies):

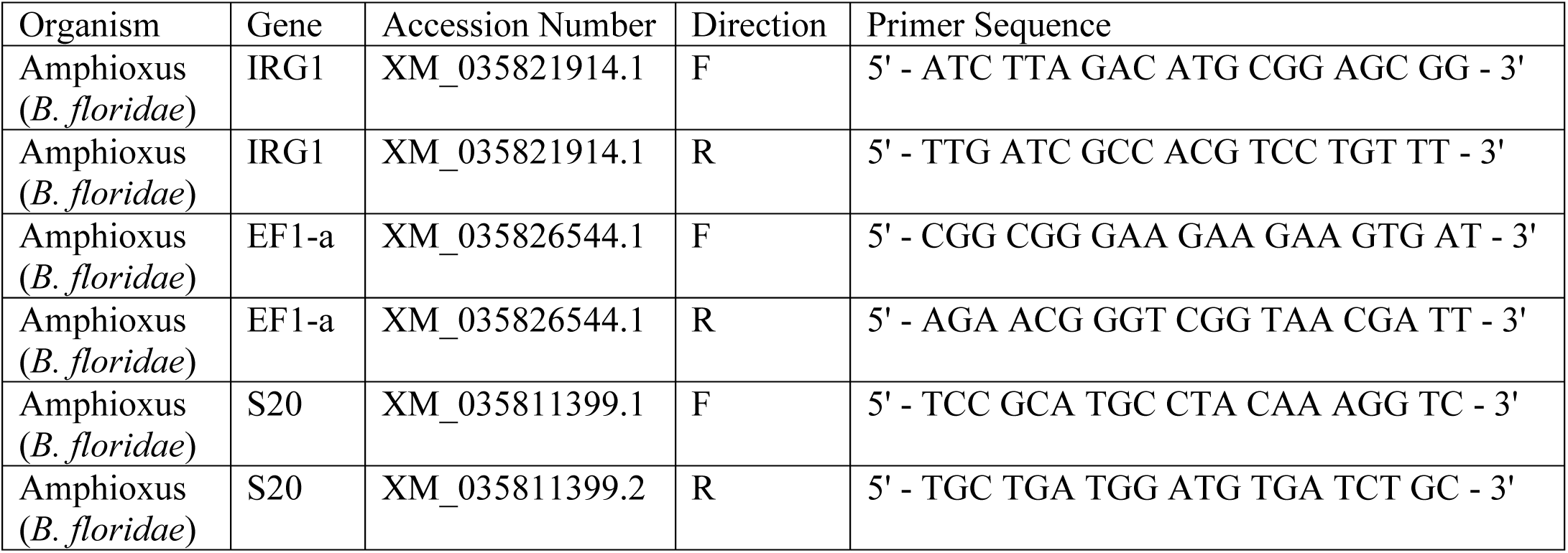

### Statistics and reproducibility

For gene expression experiments, data are shown as mean ± s.d., n ≥ 3 biologically independent samples unless stated otherwise. Statistical significance was calculated using two-tailed t test. Significance level were indicated as **** p <0.0001, *** p < 0.001, ** p < 0.01, * p < 0.05 and n.s. p > 0.05. Statistical analysis was performed using GraphPad Prism 9.3.1, or as reported by the relevant computational tools.

## Data availability

The data that support this study are available from the corresponding author upon reasonable request. The sequences used to for phylogenetic inference (.FASTA files) are available within the supplementary data.

## Author Contributions

H.S. and G.W. conceived the initial idea. R.S., A.S., and M.D. performed bioinformatics analysis; H.S., R.S., and F.M. performed recombinant protein production and enzymatic assays; R.S. and H.S. performed dissections and qRT-PCR assays. H.S. and R.S. wrote the initial manuscript with input from G.W. All authors reviewed and approved the manuscript.

## Acknowledgements

We thank Yale School of Medicine, Yale West Campus, and Yale Systems Biology Institute for the instrumentation support; X. Shi for technical support on the LC-MS; J. Liu for technical support; A. Chavan and L. Wu on advise on assays in amphioxus and oyster; K. Byrne for her insight in sequence analysis; and members of the Shen lab, D. Stadtmauer in the Wagner lab, M. Murphy, J. MacMicking, A. Goodman and J. Galán for fruitful discussion and feedback. The work is supported by Yale School of Medicine startup fund and Yale Systems Biology Institute pilot award, John Templeton Foundation to G.W. (#61329), Klingenstein-Simons fellowship Awards in Neuroscience, 1907 Foundation Trailblazer Award, and Rita Allen Foundation to H.S. and a National Science Foundation Graduate Research Fellowship to R.S. (# DGE-2036201).

## Supplementary Figures

**Supplementary figure 1.**
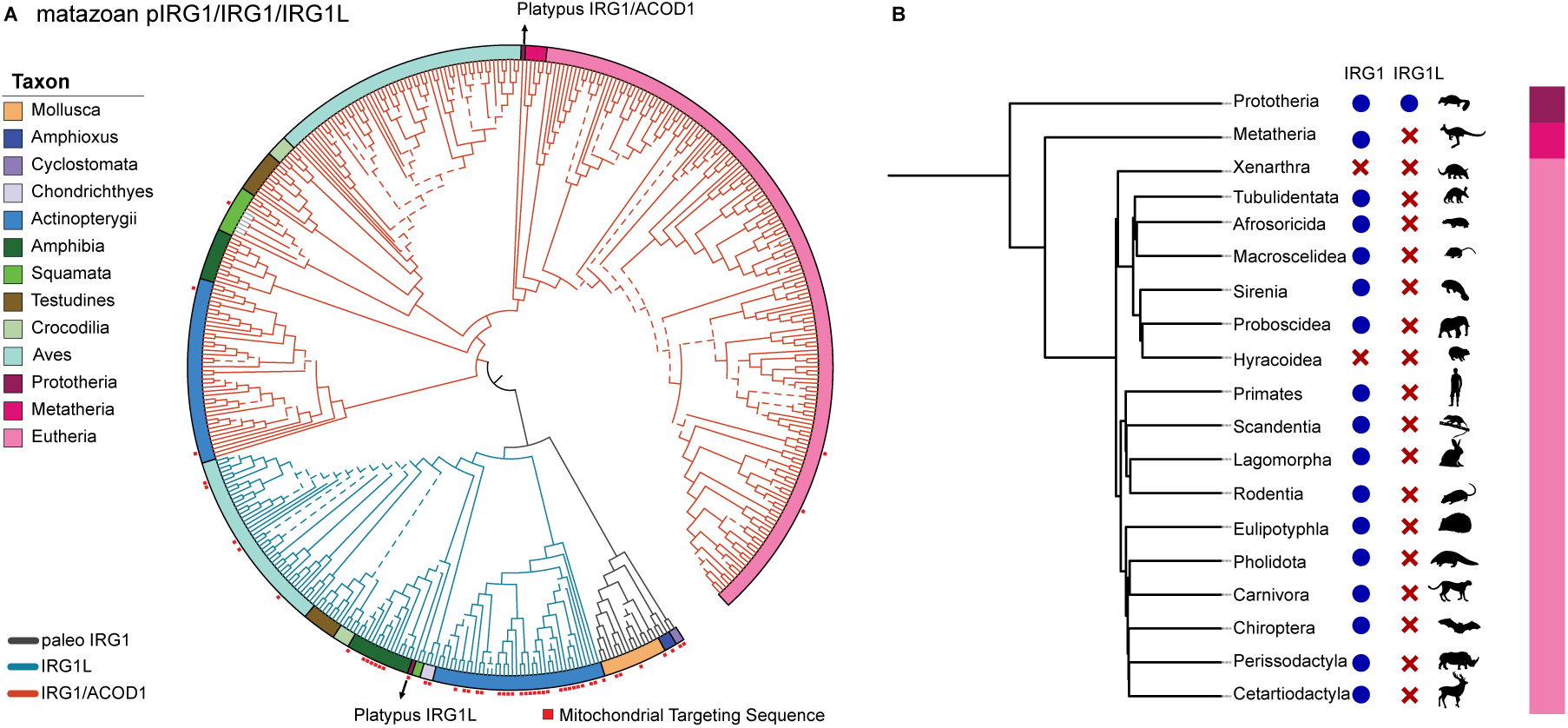
Phylogenetic tree and taxonomic distribution of pIRG1/IRG1L/IRG1 sequences in metazoa. (A) A phylogenetic tree of all metazoan IRG1 sequences. Tree branches were color-coded by three lineages: pIRG1 (black), IRG1L (cyan) and IRG1 (red). Branches with bootstrapping values higher than 70 are highlighted by solid line. Taxa are color-coded on the outer rim of the tree. The sequences with predicted Mitochondrial Targeting Sequence (MTS) were highlighted by red dots. (B) The presence of IRG1 and IRG1L in major mammalian taxa.

**Supplementary Figure 2.**
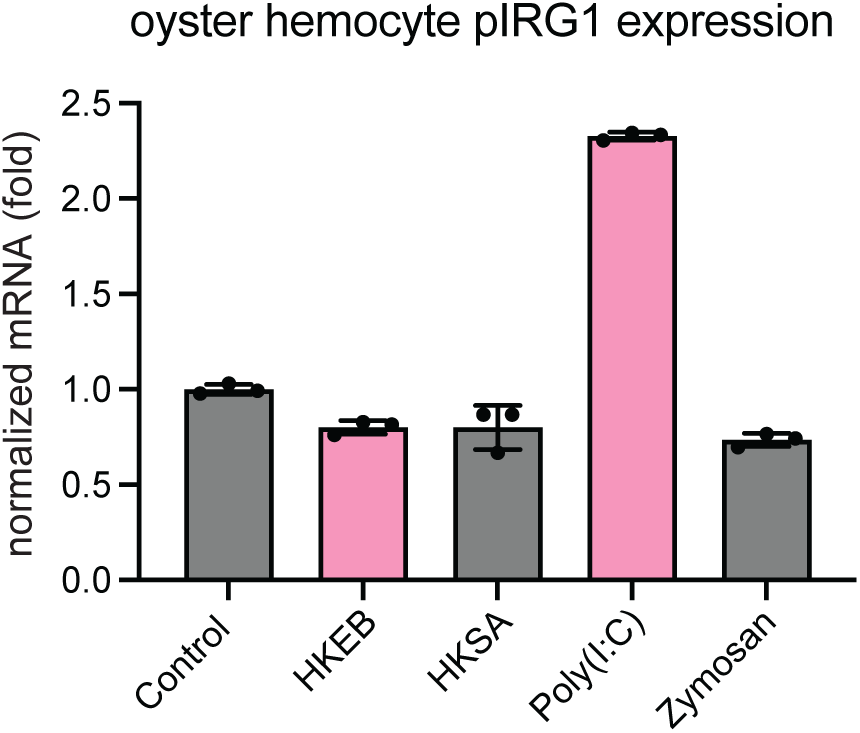
The pIRG1 is induced in the eastern oyster hemocytes by innate immune stimuli. Quantitative PCR showing the IRG1 mRNA level in hemocytes upon 8 hr stimulation of heat-killed *E. coli* 0111:B4 (HKEB) (1x10^7^ PFU), heat-killed *S. aureus* (HKSA) (1x10^7^ PFU), zymosan (100 µg/mL) and Poly (I:C) (10 µg/mL). IRG1 is significantly upregulated upon Poly(I:C) stimulation relative to other conditions.

**Supplementary File 1 The sequences used to for phylogenetic inference (.FASTA files)**

